# Habitat deterioration despite protection: long-term declines of littoral area of fishponds in Czech nature reserves

**DOI:** 10.1101/2021.12.22.473897

**Authors:** Vojtech Kolar, Kateřina Francová, Jaroslav Vrba, Stanislav Grill, David S. Boukal

## Abstract

Fishponds play a key role in current pondscapes in many developed countries. Their littoral areas, supporting multiple ecosystem functions including the maintenance of aquatic and riparian biodiversity, have been adversely affected by the move shift towards more intensive aquaculture and widespread eutrophication in the middle 20^th^ century. To counteract these changes, many fishponds received some protection, but its long-term efficiency has not been studied. Here we focus on the role of conservation status in protecting the area of littoral areas of fishponds in Czechia between the years 1950 and 2019. We found that the conservation status of these fishponds did not prevent habitat deterioration in most of the fishponds, especially during the second half of the 20^th^ century. Moreover, we detected no significant effects of the reserve establishment year, fishpond area and conservation target on the littoral areas. This suggests that the conservation measures are insufficient across fishpond reserve types. We attribute the negative trends to persisting high fish stocks, especially of common carp, and eutrophication resulting from additional feeding, pond manuring, and ongoing nutrient inputs from the pond catchments. Sediment dredging and high grazing pressure by waterfowl in some reserves can further aggravate the situation. We conclude that effective protection of the littoral areas requires a paradigm shift towards less intensive fish stock management, more frequent summer drainage, and effective reduction of all nutrient inputs to increase the water quality. Such measures can help recover the littoral areas and the associated biota.

**Graphical abstract:** 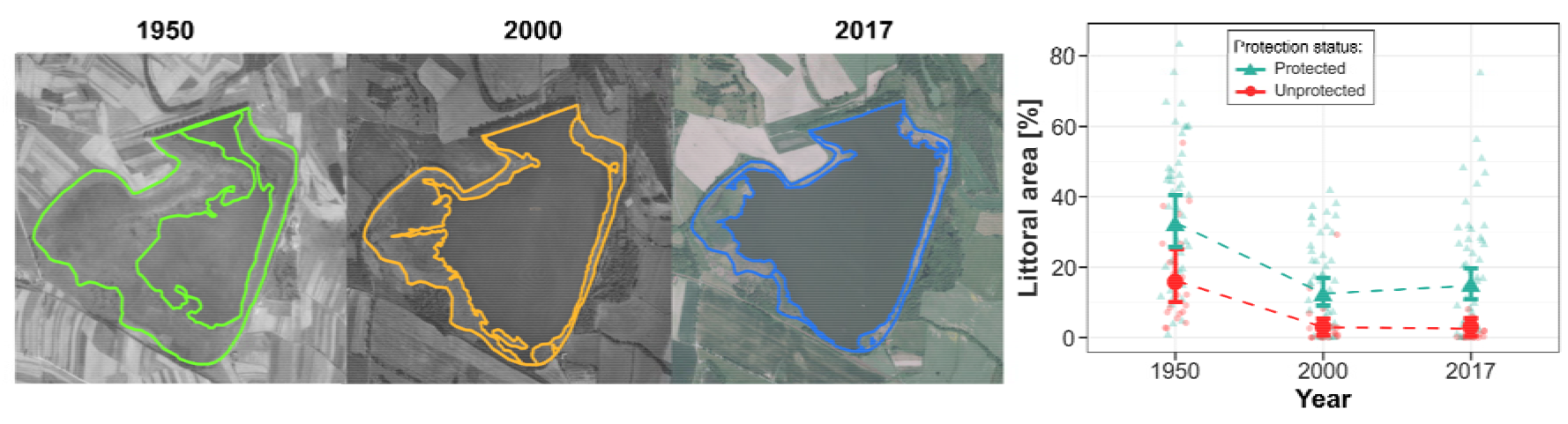

**Highlights:**

- Littoral vegetation in ponds supports high biodiversity but often lacks protection.
- We evaluated long-term changes in the littoral areas of fishponds in Czechia.
- The areas decreased markedly in both protected and unprotected ponds since 1950.
- Reserve duration, fishpond area and conservation target did not affect the trends.
- Changes in fishpond management are recommended for littoral area recovery.

## 1. Introduction

Fishponds are the most common habitat of the central European pondscapes. Most of them were created at natural or seminatural sites such as peat bogs, floodplain forests, and wetland meadows centuries ago (Francová et al., 2019). While they primarily serve for fish production and water retention, they have also become important biodiversity hotspots after deterioration and destruction of many natural wetlands (Davidson, 2014; Reid et al., 2019, Kolar and Boukal, 2020). Environmental characteristics including the extent and diversity of submerged and emergent vegetation vary immensely both within and among fishponds, which makes them ideal to support high regional biodiversity (Wezel et al., 2014). Consequently, some fishponds became protected by national and European law.

Traditional fishpond management in central Europe including fish stocking, supplementary feeding, manuring, liming, drainage, and dredging intensified heavily during the 20^th^ century to increase fish production. Moreover, phosphorus and nitrogen inputs in fishponds increased 40 and 9 times, respectively (Potužák et al., 2007). These changes led to an overall decline in the diversity of macrophytes (Francová et al., 2019), aquatic beetles (Kolar and Boukal, 2020), dragonflies (Šigutová et al., 2015), macrozoobenthos and zooplankton (Nieoczym and Kloskowski, 2014), amphibians (Kloskowski, 2010) and birds (Broyer and Curtet, 2012). Moreover, high stocks of common carp suppress large zooplankton, leading to a classical trophic cascade with frequent algal blooms and reduced water transparency (Matsuzaki et al., 2007; Potužák et al., 2007), and disturb the sediment that further reduces transparency (Roberts et al., 1995) and causes mechanical damage to submerged macrophytes (Broyer and Curtet, 2012; Matsuzaki et al., 2007). All these effects often lead to a substantial retreat of littoral areas.

Littoral areas are important hotspots of biodiversity in the fishpond ecosystems (Batzer and Wissinger, 1996; Francová et al., 2019), as most species inhabit shallow parts with submerged and emerged vegetation rather than open water (Kloskowski et al., 2020). Some taxa use these structured habitats as a refuge against predators (Warfe and Barmuta, 2004), while some predators seek littorals due to high prey availability (Kloskowski et al., 2020) or use the habitat structure to facilitate their hunting (Klecka and Boukal, 2012). Furthermore, littoral macrophytes can improve water quality, stabilize shores, and provide food for many organisms (Batzer and Wissinger, 1996; Roberts et al., 1995). Thus, their effective protection is necessary for many aquatic and semi-aquatic species, especially considering recent massive species extinctions and population decline (Eichenberg et al., 2021; Sánchez-Bayo and Wyckhuys, 2019).

While overall eutrophication and fish overstocking are known to severely affect the littoral environment, we need to develop efficient strategies to mitigate their adverse effects on pond ecosystems (Francová et al., 2019). In particular, we lack comparative studies of long-term trends of environmental conditions in fishponds and the efficiency of conservation measures such as legal protection of the fishpond habitats. To fill this gap, we analysed long-term trends in the extent of emerged macrophyte vegetation (hereafter ‘littoral area’) in legally protected and unprotected fishponds in Czechia. We expected the littoral areas in protected fishponds to remain stable or increase over time, especially in fishponds that were declared as nature reserves earlier, had a larger littoral area, or are smaller. The latter are used as nursery ponds and are usually less intensively managed than the larger main/ongrowing ponds (Francová et al., 2019). Furthermore, we hypothesized that pond reserves focused on macrophyte conservation will have larger littoral areas than those with other conservation targets (i.e., wetland communities and animals). Finally, we expected to observe declines of the total littoral areas in unprotected ponds, especially if they were large and had initially large littoral areas that were destroyed by the intensive management practices summarized above. We use our findings to highlight current issues arising in effective protection of fishpond littoral areas and propose measures required to amend the situation.

## 2. Materials and Methods

### 2.1 Study area

We analysed the effect of conservation status on the littoral areas of fishponds in South Bohemia (Fig. 1). For the sake of this study, we define fishpond as a man-made standing water body larger than 0.5 ha. More than 7,000 (total area >30,000 ha) fishponds in this region also include over 60 fishponds with a legal protection status (hereafter ‘fishpond reserves’).

**Fig. 1.**
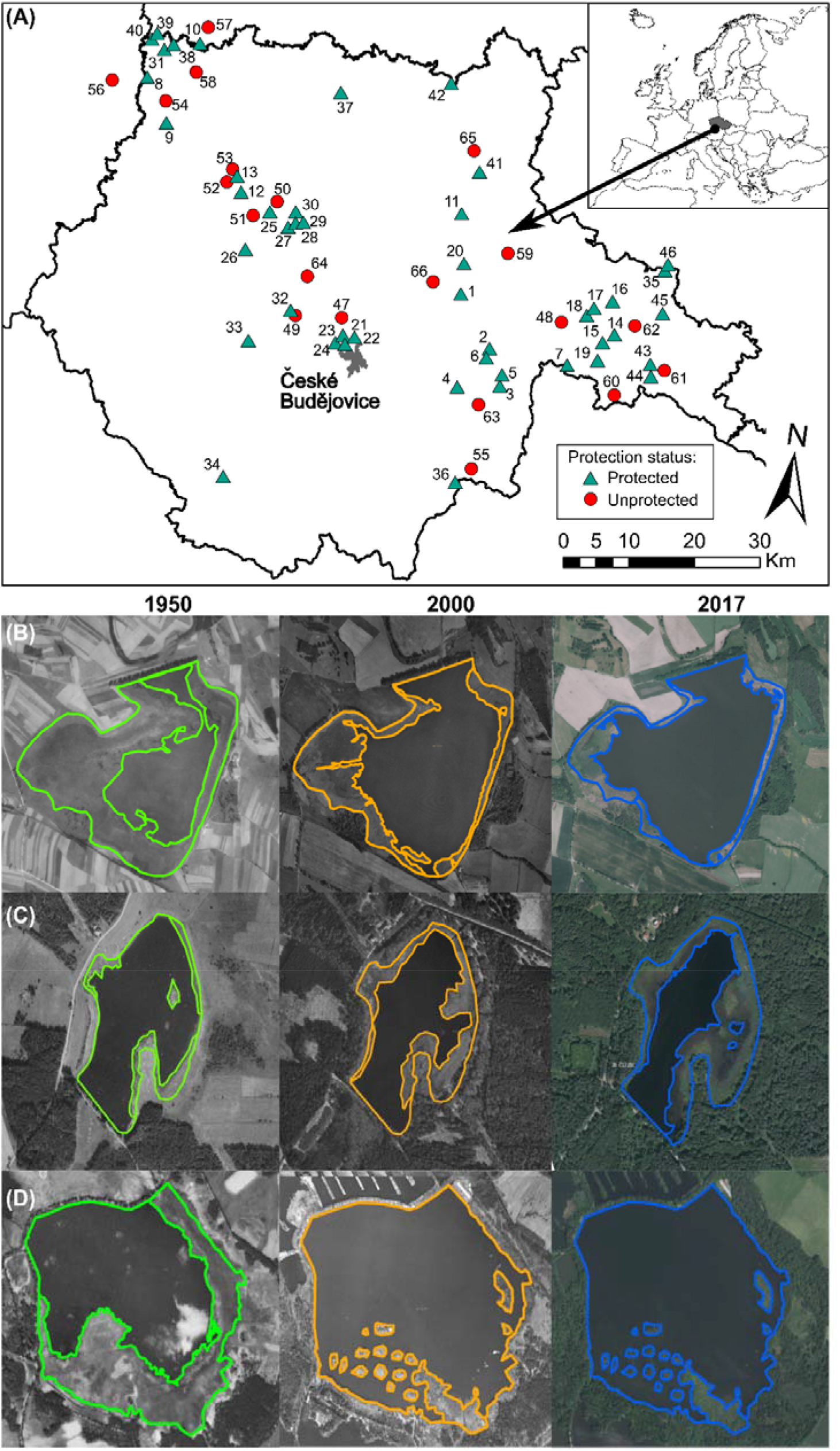
Study area and examples of temporal changes in the littoral areas of fishponds. (A) Map of South Bohemia with the studied fishponds. (B–D) comparison of three different fishponds between years 1950 (green), 2000 (orange), and 2017 (blue): decrease of the littoral area in the Řežabinec fishpond (B), increase of the littoral area in the Blanko fishpond (C), and example of dredging that resulted in artificial islands in the Staré Jezero fishpond (D). Orthophoto maps of each fishpond are on a different scale. See Table A1 for details.

### 2.2 Analyses

To quantify changes in the littoral area (i.e., the extent of the surface covered with emergent macrophyte vegetation) of fishponds, we used orthophoto maps (CUZK, 2021) from the years 1949–1953 (hereafter referred to as ‘1950’), 1998–2001 (as ‘2000’), and 2017–2019 (as ‘2017’, Table A1). We chose current fishpond reserves and located them first on the orthophoto maps from 1950; note that only three fishponds were already protected at that time (Table A1). We selected only fishponds for which high-quality images were available, and eliminated those with image noise (e.g., clouds over the fishpond) and those with unclear limits of the littoral and open-water areas. We could not locate several protected fishponds on the 1950 maps as they were likely built or restored later. This procedure yielded 46 protected and 20 unprotected fishponds; the latter were randomly selected within a radius of 20 km around a subset of the protected ones as controls (Fig. 1A).

Orthophoto maps were analysed by one person (V.K.) to avoid personal biases. All areas estimated from the maps were processed and vectorized in ArcGIS software (v. 10.7.1, ESRI, 2011). We initially estimated the total surface area of each fishpond, excluding any islands if present but including the littoral area, in the most recent maps from 2017. We then estimated the total littoral area around the shores and islands of each fishpond in the same year and years 1950 and 2000, and corrected for changes in the total surface area between the earlier dates and 2017 if necessary (Fig. 1B–D and Table A1). Littoral area was expressed as a proportion of the total fishpond area in that year. We collected the following explanatory variables for each fishpond (only for protected ones): year of reserve establishment, total surface area in 2017, littoral area in 1950, and conservation target divided into three categories: macrophytes, wetland communities (i.e., animals including amphibians, birds, insects and/or macrophytes together), and animals (amphibians and/or birds) according to the official database of the Nature Conservation Agency of the Czech Republic (AOPK ČR, 2020).

We used a model selection approach in which we built a set of candidate models and used the corrected Akaike information criterion (AICc) to rank these models and identify the most parsimonious ones (Burnham and Anderson, 2002). Temporal differences in the littoral area proportions were first analysed with generalised linear mixed models (GLMMs) using beta regression with a logit link function. Zero proportions were adjusted to 0.0001 (*n* = 11) prior to analysis. We first considered year as an explanatory variable with three levels (‘1950’, ‘2000’ and ‘2017’) and then the actual year of orthophoto imaging as a continuous explanatory variable. In both analyses, we also considered protection status as another explanatory variable and used fishpond identity as a random intercept and slope (Table A1). In the latter analysis, we considered year as a linear term or second-order polynomial and added a random intercept for the year effect. This yielded five and seven candidate models in each respective analysis (Table A2A).

We further calculated the ln-transformed ratio of the total littoral area in each fishpond reserve in 2017 and 1950 to measure its rate of change, with negative values corresponding to declining littoral areas. The data were approximately normally distributed with four strong outliers corresponding to nearly lost littoral areas. We fitted them with a set of eight robust linear regressions linking the rate of change to the conservation target (always included in the models; its significance in the most parsimonious model was assessed by an ANOVA) and a linear combination of up to three additional explanatory variables: year of establishment, total littoral area in 1950, and log10-transformed total fishpond area (Table A2C).

We used the function *glmmTMB* (Brooks et al., 2017) to analyse the GLMMs, the function *rlm* (Venables and Ripley, 2002) for the robust regressions, and function *emmeans* (Lenth, 2021) for planned comparisons in the most parsimonious model (Table A3 for details). All analyses were conducted in R (R Core Team, 2020).

## 3. Results and Discussion

### 3.1 Shrinking of littoral areas in fishponds: main patterns and likely causes

Model selection for the temporal differences with year as a factor showed that the proportion of littoral area depended on the interaction of time and protection status (Fig. 2A and Table A2). Planned comparisons confirmed that (future) reserves had more developed littoral areas than unprotected fishponds both in 1950 and in 2000 and 2017, and that littoral areas in 2000 and 2017 were significantly smaller than in 1950 in both fishpond categories (Table A3). As a result, littoral areas of the protected fishponds are now comparable to their extent in unprotected fishponds in 1950 (Fig. 2A).

**Fig. 2.**
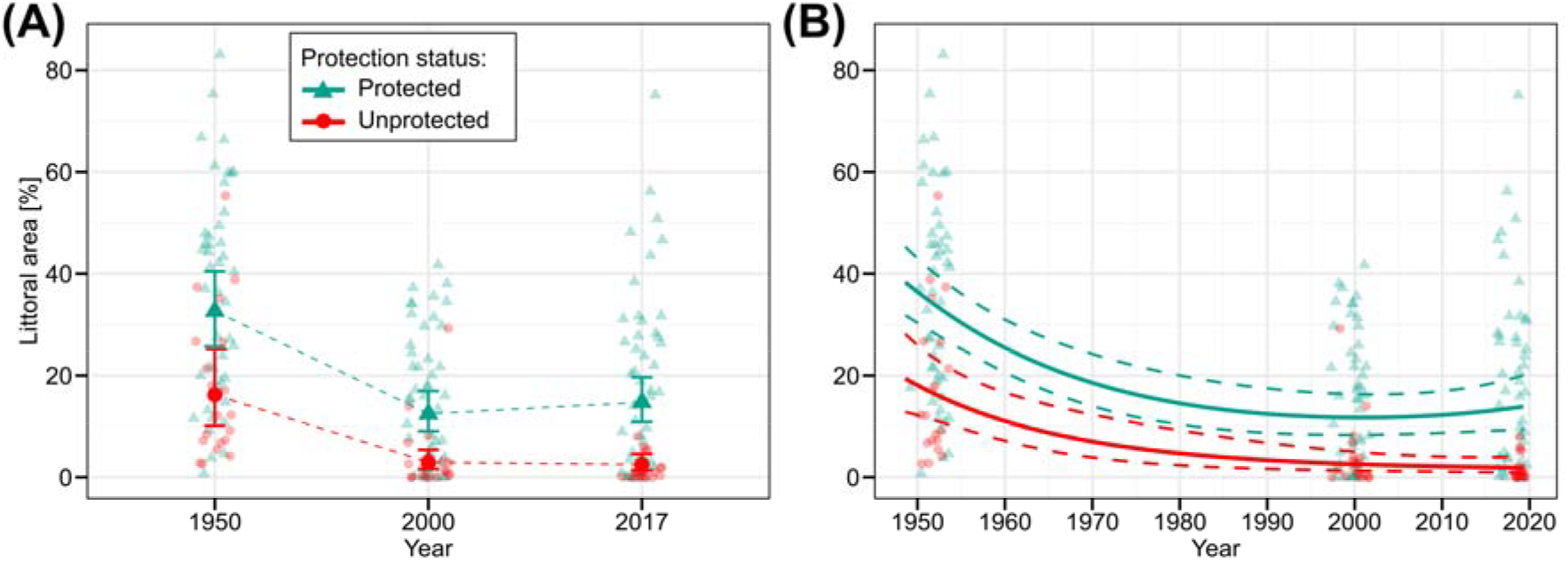
Temporal trends in the proportion of littoral area (as % of total fishpond surface area) in fishponds between 1950, 2000 and 2017. (A) changes within each fishpond category; larger symbols show mean values and error bars 95% confidence intervals. (B) overall trend predicted by the most parsimonious model; dashed lines show 95% confidence intervals. Data for individual fishponds shown as small symbols.

The most parsimonious model for the continuous temporal trends identified strong declines in the extent of littoral areas in both protected and unprotected fishponds until ca. year 2000 with a weak trend towards subsequent recovery in protected but not in unprotected fishponds, in which the decline continued (Fig. 2B). Indeed, decreases in littoral areas in protected fishponds were more common (*n*=38) and on average more dramatic (mean: −22.5%, range: −0.5 to −58.2%) than increases (*n*=8, mean: 12.5%, range: 0.8–43.0%), while littoral areas declined in all unprotected fishponds (*n*=20, mean: −16.0%, range: −2.4 to −54.0%; Table A1).

The observed degradation of littoral areas is likely connected to the intensification of fishpond management since the 1950s. Increased manuring and supplementary feeding, overstocking dominated by common carp, and spread of invasive species such as topmouth gudgeon and Prussian carp led to decreased water quality (Potužák et al., 2007). The resulting eutrophic to hypertrophic conditions cause frequent long periods of turbid water and light limitation for macrophytes, further aggravated by bioturbation and grazing on macrophytes by common carp and other cyprinids. Moreover, summer drainage can help restore littoral areas and increase the diversity of macrophytes (Šumberová et al., 2006; Broyer and Curtet, 2012; Francová et al., 2021) but is often omitted in intensive management.

Degradation or complete disappearance of some littorals could be attributed to dredging (Fig. 1D). Until recently, many fishponds were either completely or partially dredged with heavy machinery (IUCN, 1997) to increase the open water area and volume to accommodate larger fish stocks. The dredged material was often left around the banks and sometimes used to build islands providing additional bird habitats (IUCN, 1997). However, the material left on the banks devastated an important water-land transitional ecotone in many reserves and disabled littoral renewal. Widespread lack of data on the extent and timing of dredging after 1950 prevented us from quantifying its effect.

Traditional management also includes direct mowing of macrophytes (Francová et al., 2019). This practice should be strictly prohibited in fishpond reserves unless it protects focal protected species from excessive macrophyte growth and establishment of invasive species.

Finally, high waterfowl densities can degrade and destroy the littoral vegetation even in shallow parts that are inaccessible to fish. They can do so directly by grazing (e.g., geese and swans) and indirectly by release of additional nutrients (Hoyer and Canfield, 1994; Noordhuis et al., 2002).

### 3.2 Effects of fishpond area, conservation period and target conservation category on the relative change of littoral area

The most parsimonious model suggests that the relative change in the littoral area between 1950 and 2017 was unaffected by the fishpond area, littoral area in 1950, year of reserve establishment (Table A2, Fig 1A), and target conservation category (ANOVA: *F*_2_ = 0.345, *P* = 0.71). The other plausible model with fishpond area as another predictor (Table A2) showed that larger fishponds tended to more pronounced declines in the littoral area than smaller ones (details not shown).

The missing relationship between the reserve age and littoral area recovery is surprising. Our data show that neither >50 years of a reserve status may protect the littoral areas, nor up to 30 years of protection may enable recovery (see Table A1 for details). This is likely due to inappropriate management of protected fishponds. Similar negative temporal trends are known in terrestrial biota (Gray et al., 2016a; Rada et al., 2019). Site-specific conservation management for each reserve is thus needed (Gray et al. 2016), especially for fishponds that are strongly affected by both target conservation management and human impacts on the surrounding landscape and catchment (Wezel et al., 2013).

Littoral areas tended to retreat more in larger fishponds, while all cases in which the littoral area increased between 1950 and 2017 were confined to smaller ponds (surface area <10 ha). This is most likely related to fishpond management types: while smaller fishponds are usually used for fish fry breeding, larger ones are used as main ongrowing ponds (Francová et al., 2019, 2021). This does not mean that larger reserves are less valuable but highlights that they deserve proper conservation and less intensive fishery management. We thus recommend protecting fishponds with well-developed littoral areas and harbouring protected species irrespectively of the fishpond size (Francová et al. 2021; Kolář & Boukal 2020).

Interestingly, we found no link between the changes in the littoral area and the conservation target category. The trends were most variable in reserves with wetland communities as the declared conservation target and least variable (and always negative) in reserves focused on macrophytes. This intriguing result suggests a systemic failure of nature conservation across fishponds with different conservation focus. More detailed studies in these reserves should confirm whether our, admittedly coarse, measures of the ongoing changes in the littoral areas also signal changes in population sizes or species richness of the target groups.

## 4. Conclusions

Our study shows that legal protection did not avert a widespread decline of littoral areas in fishponds in Czechia since mid 20th century. This trend is detrimental as littoral areas support biodiversity across pondscapes (Vanacker et al., 2018). The observed declines signal the need for conservation agencies to balance fishpond management, focused on other services such as fish production, with conservation needs. We focused on littoral areas as indicators of overall habitat availability for various biota; further studies are required as many taxa inhabiting the littoral areas differ in their environmental requirements such as the extent or species composition of the submerged and emerged macrophytes. Requirements of different taxa could even be antagonistic (Broyer and Curtet, 2012), making general recommendations difficult. Another challenge lies in the implementation of desirable changes in fishpond management. Most Czech fishponds (including reserves) are privately owned, which limits options for externally imposed management plans.

While not a direct focus of our study, we reiterate that effective protection of the littoral part of fishpond ecosystems requires several well-known measures (Broyer and Curtet, 2012; Kloskowski, 2010): (i) use of optimal fish stock size tailored for local abiotic conditions, (ii) use of fish polyculture with a higher proportion of predatory species, (iii) more effective or no use of supplementary feed, and (iv) no manure application. Wherever possible, this should be accompanied by reducing nutrient inputs from the surrounding catchment. In addition, more frequent use of summer drainage could support the littoral area development. Taken together, these measures can reduce the nutrient loads and support growth of submerged and emerged macrophytes in the predominantly eutrophic central European pondscape.

## Acknowledgements

We acknowledge financial support by the program of the Strategy AV 21 (VP21) from the Czech Academy of Sciences. V.K. was further supported by the Grant Agency of the University of South Bohemia (GAJU 116/2019/P).

## Author statement

Vojtech Kolar: Conceptualization; Methodology; Data curation; Investigation; Visualization;

Writing – Original Draft

Kateřina Francová: Writing – Original Draft

Jaroslav Vrba: Writing – Review & Editing

Stanislav Grill: Data curation

David S. Boukal: Methodology; Validation; Investigation, Writing – Review & Editing

