## Supplementary material for "Habitat deterioration despite protection: long-term declines of littoral area of fishponds in Czech nature reserves"

**Table A1.** List of the studied fishponds. ID = number in Fig. 1; fishpond = official name of the reserve or unprotected fishpond; GPS coordinates given in WGS-84 format. Protected = protection status; established = year of the reserve establishment year; conservation target: A = animals, M = macrophytes, WA = wetland communities. Year 1950 (2000, 2017) = actual year of imaging; TA = total area in given year, LA = littoral area (both in m^2^) and relative littoral area (in %) at each time point. Abbreviations in names of protected fishponds correspond to different levels of protection: NPP = National Nature Monument, NPR = National Nature Reserve; PP = Nature Monument, PR = Nature Reserve.

| **ID** | **Fishpond** | **GPS N** | **GPS E** | **Protected** | **Established** | **Conservation target** | **Year 1950** | **TA 1950** | **LA 1950** | **Year 2000** | **TA 2000** | **LA 2000** | **Year 2017** | **TA 2017** | **LA 2017** | **PLA 1950 (%)** | **PLA 2000 (%)** | **PLA 2017 (%)** |
| --- | --- | --- | --- | --- | --- | --- | --- | --- | --- | --- | --- | --- | --- | --- | --- | --- | --- | --- |
| **1** | PR Rod | 49.12153 | 14.74554 | Yes | 1990 | A | 1953 | 317274 | 146089 | 1998 | 311729 | 92559 | 2017 | 313083 | 87065 | 46.05 | 29.69 | 27.81 |
| **2** | PR Rybníky u Vitmanova - Ženich | 49.02082 | 14.84159 | Yes | 1994 | WA | 1953 | 677611 | 406852 | 1998 | 739160 | 19841 | 2017 | 731561 | 13838 | 60.04 | 2.68 | 1.89 |
| **3** | NPP Vizír | 48.96304 | 14.88656 | Yes | 1987 | M | 1952 | 96846 | 72958 | 1998 | 84307 | 8845 | 2017 | 78580 | 34266 | 75.33 | 10.49 | 43.61 |
| **4** | PR Ruda u Kojákovic | 48.94692 | 14.77095 | Yes | 1991 | WA | 1952 | 770611 | 334224 | 1998 | 675987 | 146052 | 2017 | 683462 | 193553 | 43.37 | 21.61 | 28.32 |
| **5** | PR Staré jezero | 48.97836 | 14.88991 | Yes | 1994 | WA | 1952 | 882729 | 356595 | 1998 | 847654 | 41376 | 2017 | 847654 | 41376 | 40.40 | 4.88 | 4.88 |
| **6** | NPR Nová a Stará řeka - Nový Spálený rybník | 49.00782 | 14.83927 | Yes | 1956 | WA | 1953 | 145493 | 86866 | 1998 | 145493 | 50268 | 2017 | 142851 | 68884 | 59.70 | 34.55 | 48.22 |
| **7** | PP Blanko | 49.01291 | 15.06954 | Yes | 1998 | WA | 1952 | 84308 | 11127 | 2000 | 84308 | 29993 | 2017 | 84030 | 47259 | 13.20 | 35.58 | 56.24 |
| **8** | PP Nový rybník U Lnář | 49.4373 | 13.78309 | Yes | 1933 | A | 1951 | 282069 | 1987 | 2001 | 298890 | 602 | 2017 | 298890 | 602 | 0.70 | 0.20 | 0.20 |
| **9** | PR Kovašinské louky Kovašín | 49.3586 | 13.85261 | Yes | 1990 | WA | 1951 | 89755 | 51987 | 2001 | 91451 | 7510 | 2017 | 86256 | 1530 | 57.92 | 8.21 | 1.77 |
| **10** | PP Drážská Koupě | 49.51551 | 13.91309 | Yes | 2014 | A | 1952 | 55450 | 14327 | 2001 | 56510 | 12312 | 2017 | 51244 | 13639 | 25.84 | 21.79 | 26.62 |
| **11** | PP Nový rybník u Soběslavy | 49.26635 | 14.72098 | Yes | 1949 | WA | 1953 | 129271 | 57765 | 1999 | 130174 | 48546 | 2017 | 130099 | 41250 | 44.68 | 37.29 | 31.71 |
| **12** | NPR Režabinec | 49.25343 | 14.0921 | Yes | 1949 | A | 1951 | 937080 | 621943 | 2000 | 937014 | 301311 | 2017 | 937080 | 190172 | 66.37 | 32.16 | 20.29 |
| **13** | PP Velký Potočný | 49.27768 | 14.06425 | Yes | 1985 | WA | 1951 | 393369 | 129303 | 2000 | 393369 | 71096 | 2017 | 393369 | 66887 | 32.87 | 18.07 | 17.00 |
| **14** | PR Skalák u Senotína | 49.06364 | 15.15839 | Yes | 2002 | WA | 1953 | 39378 | 16249 | 2000 | 39141 | 5996 | 2017 | 39139 | 18267 | 41.26 | 15.32 | 46.67 |
| **15** | NPP Kaproun | 49.07969 | 15.18989 | Yes | 1987 | M | 1953 | 5792 | 2562 | 2000 | 6152 | 0.01 | 2017 | 5816 | 822 | 44.23 | 0.00 | 14.13 |
| **16** | PR Hrádeček | 49.14277 | 15.17066 | Yes | 2002 | WA | 1953 | 92814 | 4272 | 2000 | 92814 | 0.01 | 2017 | 92814 | 0.01 | 4.60 | 0.00 | 0.00 |
| **17** | NPP Krvavý rybník | 49.12508 | 15.12212 | Yes | 1994 | WA | 1953 | 1220911 | 238467 | 2000 | 1183001 | 5042 | 2017 | 1164407 | 40596 | 19.53 | 0.43 | 3.49 |
| **18** | NPP Kačležský rybník | 49.1116 | 15.10479 | Yes | 1994 | WA | 1953 | 1524378 | 177035 | 2000 | 1520744 | 8184 | 2017 | 1521526 | 60898 | 11.61 | 0.54 | 4.00 |
| **19** | EVL Osika | 49.03012 | 15.14993 | Yes | 2009 | M | 1953 | 648670 | 24879 | 2000 | 617291 | 15501 | 2017 | 630517 | 5676 | 3.84 | 2.51 | 0.90 |
| **20** | PP Farářský rybník | 49.17552 | 14.74212 | Yes | 1988 | M | 1953 | 63288 | 31305 | 1998 | 62342 | 9964 | 2017 | 63288 | 2565 | 49.46 | 15.98 | 4.05 |
| **21** | PR Vrbenské rybníky - Bažina | 49.00696 | 14.43964 | Yes | 1990 | WA | 1952 | 65021 | 38914 | 1998 | 62262 | 23716 | 2019 | 55970 | 5439 | 59.85 | 38.09 | 9.72 |
| **22** | PR Vrbenské rybníky - Starý Vrbenský | 49.0108 | 14.4334 | Yes | 1990 | WA | 1952 | 128279 | 30649 | 1998 | 128660 | 0.00 | 2019 | 128660 | 0.01 | 23.89 | 0.00 | 0.00 |
| **23** | PR Vrbenské rybníky - Nový Vrbenský | 49.00756 | 14.44314 | Yes | 1990 | WA | 1952 | 135905 | 37567 | 1998 | 121568 | 1815 | 2019 | 120606 | 0.01 | 27.64 | 1.49 | 0.00 |
| **24** | PR Vrbenské rybníky - Černiš | 49.0046 | 14.42801 | Yes | 1990 | WA | 1952 | 359741 | 67719 | 1998 | 355590 | 0.01 | 2019 | 347364 | 0.01 | 18.82 | 0.00 | 0.00 |
| **25** | PP Skalský rybník | 49.22215 | 14.18215 | Yes | 1985 | WA | 1951 | 78291 | 28459 | 2000 | 93340 | 5633 | 2019 | 88168 | 16628 | 36.35 | 6.04 | 18.86 |
| **26** | PR Záhorský rybník | 49.1473 | 14.12185 | Yes | 1997 | M | 1951 | 188623 | 70027 | 2000 | 201739 | 63428 | 2019 | 194315 | 48995 | 37.13 | 31.44 | 25.21 |
| **27** | PP Zelendárky - Starý u Krče | 49.20823 | 14.25694 | Yes | 1985 | WA | 1952 | 19249 | 5221 | 2000 | 17612 | 5237 | 2019 | 16050 | 6178 | 27.12 | 29.73 | 38.49 |
| **28** | PP Zelendárky - Nový u Krče | 49.21084 | 14.25458 | Yes | 1985 | WA | 1952 | 40858 | 8895 | 2000 | 39926 | 3050 | 2019 | 37228 | 3040 | 21.77 | 7.64 | 8.17 |
| **29** | PP Zelendárky - Na Rejčových rybník | 49.21114 | 14.26304 | Yes | 1985 | WA | 1952 | 13065 | 5978 | 2000 | 12221 | 2162 | 2019 | 12095 | 1926 | 45.76 | 17.70 | 15.92 |
| **30** | PP Zelendárky - Skopec | 49.23065 | 14.2532 | Yes | 1985 | WA | 1952 | 37051 | 7404 | 2000 | 34012 | 11605 | 2019 | 31016 | 9666 | 19.98 | 34.12 | 31.16 |
| **31** | PP Závišinský potok - Luh | 49.5022 | 13.84227 | Yes | 2013 | WA | 1952 | 57586 | 8533 | 2001 | 57586 | 2294 | 2019 | 56229 | 1682 | 14.82 | 3.98 | 2.99 |
| **32** | PP Velký Karasín | 49.04512 | 14.27931 | Yes | 1991 | WA | 1952 | 63198 | 32939 | 2000 | 62397 | 14518 | 2019 | 62397 | 14887 | 52.12 | 23.27 | 23.86 |
| **33** | PP Koubovský rybník | 48.98112 | 14.17004 | Yes | 1988 | M | 1952 | 13109 | 4515 | 2000 | 12661 | 1950 | 2019 | 10814 | 2850 | 34.44 | 15.41 | 26.36 |
| **34** | PR Pláničský rybník | 48.72286 | 14.15534 | Yes | 1996 | M | 1949 | 104178 | 18339 | 2000 | 104560 | 22750 | 2019 | 100917 | 9069 | 17.60 | 21.76 | 8.99 |
| **35** | NPP Zhejral - Karhov | 49.21237 | 15.30777 | Yes | 1982 | WA | 1953 | 206814 | 19294 | 2000 | 217167 | 240 | 2019 | 217167 | 1575 | 9.33 | 0.11 | 0.73 |
| **36** | PP Přesličkový rybník - Přesličkový rybník | 48.77089 | 14.80307 | Yes | 1991 | WA | 1952 | 15560 | 10400 | 2001 | 14980 | 3897 | 2019 | 15419 | 4886 | 66.84 | 26.01 | 31.69 |
| **37** | PP Boukal - Zlatina | 49.46157 | 14.33364 | Yes | 1985 | WA | 1952 | 24607 | 11788 | 2000 | 26451 | 4561 | 2019 | 25630 | 7875 | 47.91 | 17.24 | 30.73 |
| **38** | PP Újezdec - Pláninský rybník | 49.49503 | 13.81925 | Yes | 2013 | WA | 1951 | 50232 | 30759 | 2001 | 50082 | 20910 | 2019 | 47942 | 24397 | 61.23 | 41.75 | 50.89 |
| **39** | PP Rybník Vočert a Lazy - Lazy | 49.51693 | 13.78727 | Yes | 2012 | A | 1952 | 44882 | 20509 | 2001 | 42640 | 8580 | 2019 | 42901 | 3015 | 45.70 | 20.12 | 7.03 |
| **40** | PP Rybnik Vočert a Lazy - Vočert | 49.51969 | 13.78776 | Yes | 2012 | A | 1952 | 40126 | 8568 | 2001 | 34232 | 5502 | 2019 | 33391 | 3706 | 21.35 | 16.07 | 11.10 |
| **41** | NPP Luční - Luční | 49.37293 | 14.74311 | Yes | 1988 | M | 1953 | 69934 | 29451 | 2000 | 87838 | 2291 | 2019 | 89631 | 2225 | 42.11 | 2.61 | 2.48 |
| **42** | PP Suchdol | 49.5043 | 14.64078 | Yes | 2012 | A | 1953 | 11124 | 1198 | 2000 | 6218 | 2125 | 2019 | 8982 | 1975 | 10.77 | 34.17 | 21.98 |
| **43** | PP Dědek u Slavonic | 49.01316 | 15.30794 | Yes | 1995 | WA | 1953 | 22877 | 3589 | 2000 | 23419 | 3143 | 2019 | 23419 | 6436 | 15.69 | 13.42 | 27.48 |
| **44** | PP Velký Troubný | 49.03436 | 15.30037 | Yes | 1995 | WA | 1953 | 33441 | 15848 | 2000 | 37132 | 2608 | 2019 | 37132 | 7705 | 47.39 | 7.02 | 20.75 |
| **45** | PR Rašeliniště Radlice | 49.13269 | 15.31599 | Yes | 2011 | WA | 1953 | 6891 | 5726 | 2000 | 6891 | 1685 | 2019 | 6891 | 5178 | 83.10 | 24.46 | 75.15 |
| **46** | NPP Zhejral - Zhejral | 49.22139 | 15.3124 | Yes | 1982 | WA | 1953 | 78822 | 9139 | 2000 | 78822 | 5721 | 2019 | 78822 | 13209 | 11.59 | 7.26 | 16.76 |
| **47** | Munický rybník | 49.04694 | 14.42151 | No | - | - | 1952 | 963638 | 70525 | 1999 | 1076099 | 3332 | 2019 | 1062437 | 942 | 7.32 | 0.31 | 0.09 |
| **48** | Dřevo | 49.09482 | 15.03666 | No | - | - | 1953 | 531114 | 22301 | 2000 | 567120 | 16359 | 2019 | 573878 | 1555 | 4.20 | 2.88 | 0.27 |
| **49** | Hlásný rybník | 49.04157 | 14.29019 | No | - | - | 1952 | 174795 | 96698 | 2000 | 149977 | 5989 | 2019 | 160872 | 2118 | 55.32 | 3.99 | 1.32 |
| **50** | Selibovský rybník | 49.2442 | 14.19659 | No | - | - | 1951 | 318523 | 23004 | 2000 | 422716 | 13660 | 2019 | 412291 | 4490 | 7.22 | 3.23 | 1.09 |
| **51** | Tvrzský rybník | 49.21257 | 14.13574 | No | - | - | 1951 | 75208 | 9114 | 2000 | 80695 | 2132 | 2019 | 80227 | 724 | 12.12 | 2.64 | 0.90 |
| **52** | Novokestřanský rybník | 49.26981 | 14.04948 | No | - | - | 1951 | 74281 | 2007 | 2000 | 73036 | 788 | 2019 | 72816 | 213 | 2.70 | 1.08 | 0.29 |
| **53** | Dobevský rybník | 49.29086 | 14.0629 | No | - | - | 1951 | 289451 | 35404 | 2000 | 308671 | 25372 | 2019 | 292337 | 15523 | 12.23 | 8.22 | 5.31 |
| **54** | Ostrý rybník | 49.40219 | 13.84149 | No | - | - | 1951 | 62511 | 16689 | 2001 | 60629 | 0.00 | 2019 | 58044 | 0.00 | 26.70 | 0.00 | 0.00 |
| **55** | Nakolický rybník | 48.80361 | 14.84135 | No | - | - | 1952 | 486509 | 83254 | 2001 | 461835 | 64550 | 2019 | 474674 | 9564 | 17.11 | 13.98 | 2.01 |
| **56** | Široký rybník | 49.4248 | 13.68215 | No | - | - | 1951 | 184196 | 12439 | 2001 | 226738 | 0.00 | 2019 | 217229 | 0.00 | 6.75 | 0.00 | 0.00 |
| **57** | Bubovický rybník | 49.54732 | 13.9324 | No | - | - | 1952 | 190892 | 5359 | 2001 | 188351 | 781 | 2019 | 180540 | 0.00 | 2.81 | 0.41 | 0.00 |
| **58** | Velký Málkovský rybník | 49.46748 | 13.90636 | No | - | - | 1951 | 83309 | 32365 | 2001 | 97084 | 0.00 | 2019 | 94714 | 0.00 | 38.85 | 0.00 | 0.00 |
| **59** | Kardaš | 49.20725 | 14.86126 | No | - | - | 1953 | 583103 | 155055 | 1999 | 591412 | 172884 | 2019 | 550193 | 10044 | 26.59 | 29.23 | 1.83 |
| **60** | Návarský rybník | 48.9712 | 15.21083 | No | - | - | 1952 | 71729 | 25188 | 2000 | 67504 | 2591 | 2019 | 74967 | 4449 | 35.12 | 3.84 | 5.93 |
| **61** | Vlastkovecký rybník | 49.03328 | 15.34023 | No | - | - | 1953 | 66221 | 14145 | 2000 | 64755 | 0.00 | 2019 | 63937 | 5158 | 21.36 | 0.00 | 8.07 |
| **62** | Popelák | 49.10312 | 15.24171 | No | - | - | 1953 | 23600 | 2156 | 2000 | 22636 | 1545 | 2019 | 19476 | 0.00 | 9.13 | 6.83 | 0.00 |
| **63** | Nový u Cepu | 48.92099 | 14.83514 | No | - | - | 1952 | 168323 | 30334 | 1998 | 163974 | 0.00 | 2019 | 162668 | 8893 | 18.02 | 0.00 | 5.47 |
| **64** | Blatec | 49.1166 | 14.30907 | No | - | - | 1952 | 830600 | 70556 | 2000 | 950019 | 18689 | 2019 | 938990 | 11565 | 8.49 | 1.97 | 1.23 |
| **65** | Jezero | 49.38678 | 14.73252 | No | - | - | 1953 | 291849 | 109140 | 2000 | 374590 | 2639 | 2019 | 371434 | 4070 | 37.40 | 0.70 | 1.10 |
| **66** | Bošilecký rybník | 49.13845 | 14.66308 | No | - | - | 1953 | 1616591 | 88711 | 1998 | 1658246 | 24427 | 2019 | 1666003 | 2414 | 5.49 | 1.47 | 0.14 |

**Table A2.** Summary of model comparisons. Values of ∆AICc, degrees of freedom (df) and Akaike weights (w) of the different models for changes of littoral area in fishponds when year is treated as a factor or as a continuous variable, and for the relative change of the littoral area in fishpond reserves. We retained all plausible models and illustrated the most parsimonious models using the package *ggeffect* (v. 0.14.1; Lüdecke, 2018). The most parsimonious and other plausible models (∆AICc ≤ 2) for each response are in bold. Models ordered by the degree of parsimony. Explanatory variables: Y_F_ = year of imaging (as factor); Y = exact year of imaging (as continuous variable); A = total surface area of fishpond; C = conservation target (factor); E = year of reserve establishment; L = initial relative littoral area.

| **Response** | **Model structure** | **∆AICc** | **df** | **W** |
| --- | --- | --- | --- | --- |
| **Littoral area, year as factor** | **~ Y_F_ × Protected + (1\|locality)** | **0** | **8** | **0.850** |
|  | ~ Y**_F_** + Protected + (1\|locality) | 3.5 | 6 | 0.150 |
|  | ~ Y**_F_** + (1\|locality) | 24.8 | 5 | <0.001 |
|  | ~ Protected + (1\|locality) | 75.5 | 4 | <0.001 |
|  | ~ 1 + (1\|locality) | 95.9 | 3 | <0.001 |
| **Littoral area, continuous year** | **~ Protected × (Y + Y^2^) + (1+Y\|locality)** | **0** | **10** | **0.837** |
|  | ~ Protected + Y + Y^2^ + (1+Y\|locality) | 3.4 | 8 | 0.151 |
|  | ~ Protected **×** Y + (1+Y\|locality) | 8.4 | 8 | 0.012 |
|  | ~ Y + Y^2^ + (1+Y\|locality) | 19.4 | 7 | <0.001 |
|  | ~ Protected + (1+Y\|locality) | 67.0 | 6 | <0.001 |
|  | ~ Y + (1+Y\|locality) | 27.9 | 6 | <0.001 |
|  | ~ 1 + (1+Y\|locality) | 84.4 | 5 | <0.001 |
| **Relative change of littoral area** | **C** | **0** | **4** | **0.368** |
|  | **C + log_10_(A)** | **0.7** | **5** | **0.253** |
|  | C + L | 2.5 | 5 | 0.107 |
|  | C + E | 2.6 | 5 | 0.101 |
|  | C + E + log_10_(A) | 3.3 | 6 | 0.072 |
|  | C + log_10_(A) + L | 3.8 | 6 | 0.056 |
|  | C + E + L | 5.2 | 6 | 0.028 |
|  | C + E + L + log_10_(A) | 6.4 | 7 | 0.015 |

**Table A3**. Summary of planned comparisons for the first analysis with year as a factor. We compared differences in the littoral area between protected and unprotected fishponds in 1950 and in 2017, and differences between the state in 1950 and in 2000 and 2017 taken together; the latter were done separately for each protection status. Results given as the parameter estimate with 95% confidence interval on the predictor scale, value of the *t*-test statistics, and corresponding *P* value; df = 190 for all comparisons.

| **Contrast** | **Estimate** | ***t*** | ***P*** |
| --- | --- | --- | --- |
| Unprotected vs. Protected in 1950 | −0.92 (−1.72 to −0.12) | −2.85 | 0.018 |
| Unprotected vs. Protected in 2017 | −1.90 (−2.79 to −1.02) | −5.35 | <0.001 |
| Unprotected: 1950–2000s | 1.93 (1.26–2.61) | 7.14 | <0.001 |
| Protected: 1950–2000s | 1.13 (0.76–1.49) | 7.70 | <0.001 |

Fig. 1A. The relationship between the relative change in the littoral area and the conservation target (A), the total fishpond water area (B), the initial relative littoral area (C), and the year of establishment (D). Conservation target categories: yellow circles = animals, red triangles = macrophytes, black squares = wetland communities.

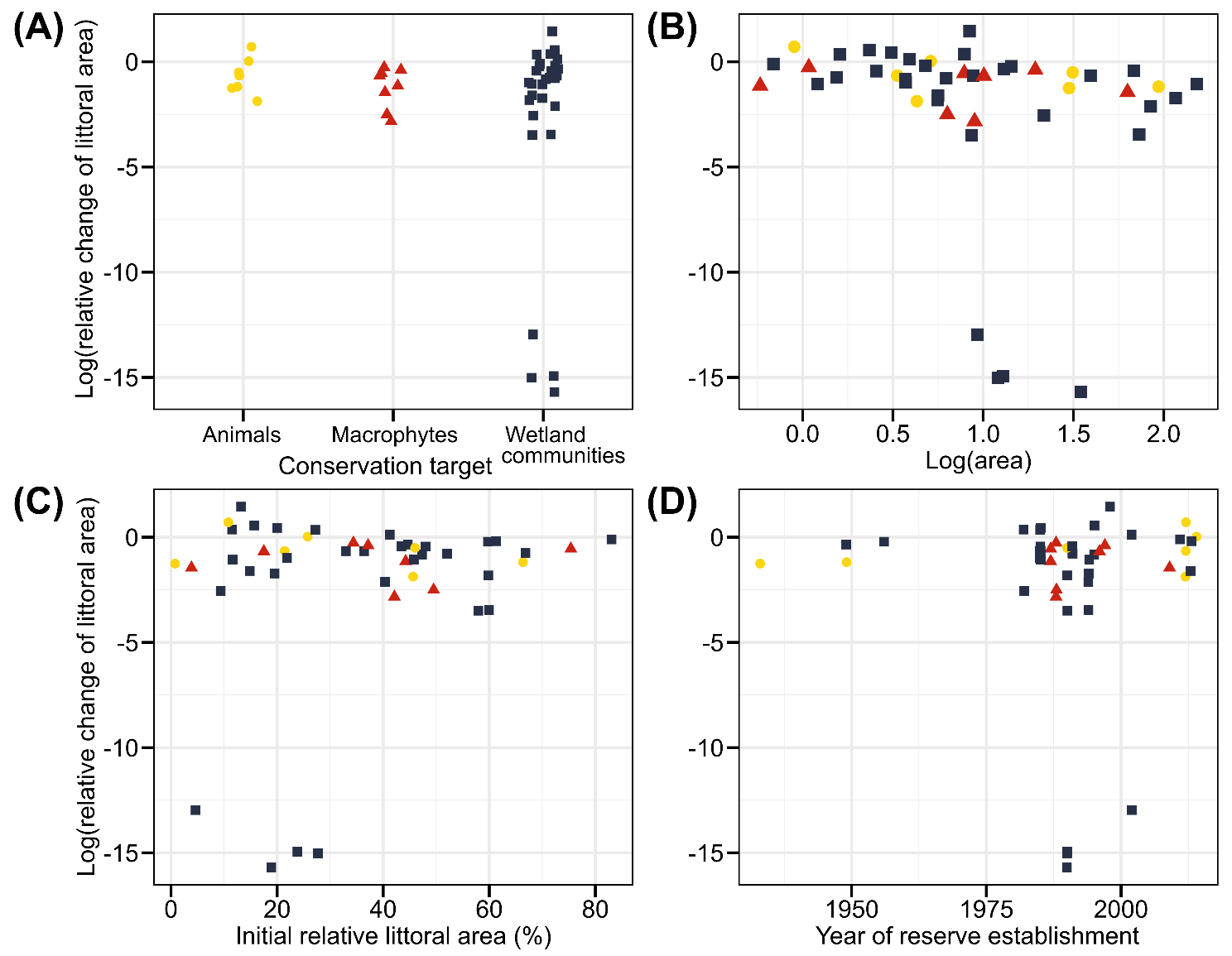
